# A cautionary note on the use of N-acetylcysteine as a reactive oxygen species antagonist to assess copper mediated cell death

**DOI:** 10.1101/2023.07.19.549632

**Authors:** Rebecca E. Graham, Richard J. R. Elliott, Alison F. Munro, Neil O. Carragher

**Affiliations:** Centre for Clinical Brain Sciences, University of Edinburgh, Edinburgh, UK; Cancer Research UK Edinburgh Centre, MRC Institute of Genetics and Cancer, University of Edinburgh, Western General Hospital, Edinburgh, UK

## Abstract

A new form of cell death has recently been proposed involving copper-induced cell death, termed cuproptosis. This new form of cell death has been widely studied in relation to a novel class of copper ionophores, including elesclomol and disulfiram. However, the exact mechanism leading to cell death remains contentious. The oldest and mostly widely accepted biological mechanism is that the accumulated intracellular copper leads to excessive build-up of reactive oxygen species and that this is what ultimately leads to cell death. Most of this evidence is largely based on studies using N-acetylcysteine (NAC), an antioxidant, to relieve the oxidative stress and prevent cell death. However, here we have demonstrated using inductively coupled mass-spectrometry, that NAC pretreatment significantly reduces intracellular copper uptake triggered by the ionophores, elesclomol and disulfiram, suggesting that reduction in copper uptake, rather than the antioxidant activity of NAC, is responsible for the diminished cell death. We present further data showing that key mediators of reactive oxygen species are not upregulated in response to elesclomol treatment, and further that sensitivity of cancer cell lines to reactive oxygen species does not correlate with sensitivity to these copper ionophores. Our findings are in line with several recent studies proposing the mechanism of cuproptosis is instead via copper mediated aggregation of proteins, resulting in proteotoxic stress leading to cell death.

Overall, it is vital to disseminate this key piece of information regarding NAC’s activity on copper uptake since new research attributing the effect of NAC on copper ionophore activity to quenching of reactive oxygen species is being published regularly and our studies suggest their conclusions may be misleading.

## Introduction

It has recently been revealed that a novel class of copper ionophores display highly selective and potent cytotoxic activity towards cancer cells. These compounds bind free copper in the media and act as a shuttle, bringing it into cancer cells selectively where it accumulates and induces cytotoxic activity [1–4].

Structurally this class of copper ionophores, including elesclomol and disulfiram contain thiocarbonyl groups, and they are known copper chelators. Disulfiram, a thiuram disulphide, breaks down in acidic or Cu(II) rich environments to produce a dithiocarbamate; diethyldithiocarbamate (DDTC) [5]. DDTC, like other dithiocarbamates, are known to form complexes with transition elements, but are most stable in a Cu(II) chelate [6]. Although elesclomol is structurally unrelated to the dithiocarbamates, it also forms organometallic complexes, particularly with Cu(II), due to its two thiocarbonyl moieties. We and others, have previously demonstrated that elesclomol and disulfiram act as copper ionophores. Incubation of cancer cells with elesclomol and disulfiram leads to a significant increase in intracellular copper levels and eventually cell death. Critically, removal of free copper from the media prior to drug treatment results in complete loss of the drugs cytotoxic effects, demonstrating the essentiality of copper in the cytotoxic mechanism-of-action of these compounds [1–4,7].

While there is now much evidence demonstrating the role of intracellular copper accumulation in the mechanism of these compounds, the mechanism leading to the cell death downstream of the copper accumulation is still contested. Since copper is a redox active metal, it has been proposed that the intracellular copper accumulation leads to the production of reactive oxygen species (ROS) and that this is the cause of cell death [4,8,9]. Much of the evidence for this is based on the ability of N-acetylcysteine (NAC) to alleviate the toxicity of the drugs. NAC is a widely used, powerful antioxidant *in vivo* and *in vitro*. NAC is a precursor of L-cysteine that results in glutathione biosynthesis. It acts directly as a scavenger of free radicals, especially oxygen radicals. Due to the fact that NAC is a ROS antagonist, these results have been interpreted to support the claim that ROS may underlie the mechanism by which copper ionophores induce cuproptosis. However, NAC has other, less well known mechanisms, including an ability to form conjugates with copper [10] and thiol-reactive compounds [11]. Here we explore the mechanism through which NAC mediates relief of copper ionophore induced cell death and importantly demonstrate that it is not via the widely accepted effects of ROS antagonism.

## Materials and Methods

### Cell culture

Oesophageal adenocarcinoma (OAC) lines were grown in Roswell Park Memorial Institute (RPMI) (Life Technologies; #11875101) supplemented with FBS (10 %, Life Technologies; #16140071) and L-glutamine (2 mM, Life Technologies; #A2916801) and incubated under standard tissue culture conditions (37 °C and 5 % CO2). The oesophageal epithelial line EPC2-hTERT was grown in KSFM (Life Technologies; #17005075) supplemented with human recombinant epidermal growth factor (5g/L) and bovine pituitary extract (50 mg/L).

For subculture of OAC lines, cell were detached with trypsin (0.25 %, 1 mL, 5 minutes, 37 °C), and neutralised with fresh growth media. For subculture of EPC2-hTERT, soy-bean trypsin inhibitor (250 mg/L, 5 mL) was added to neutralise the trypsin and centrifuged (5 minutes, 250 x g). The cell pellet was then re-suspended in fresh media.

Cells were seeded (50 µL per well) at 1500 cells per well except SK-GT-4 which was seeded at 1000 cells per well into 384-well, CELLSTAR^®^ Cell Culture Microplates (Greiner, #781091), and incubated under standard tissue culture conditions for 24 hrs before the addition of compounds.

### N-acetylcysteine dose response

Disulfiram and elesclomol dose responses were carried in the presence and absence of a 15 min pre-incubation with 1 or 10 mM NAC (A7250; Sigma). NAC was made up in media and 0.1M NaOH was used to neutralise the media and bring the pH back to 7.4. This was then syringe filtered before being added to the cells.

### H_2_O_2_ dose response

H_2_O_2_ (H1009; Sigma) was diluted in media to give an 18 point dose response starting at 1 mM and incubated for 48 hrs. Viability was assessed by Alamar Blue.

### Inductively coupled mass spectrometry

5 × 10^6^ cells were seeded in a T175 flask and incubated in 25ml RPMI overnight. Media was then replaced with the addition of compound treatments (DMSO (0.1%), disulfiram (600 nM), or elesclomol (200 nM)) with or without a 15 min pre-incubation with 10 mM NAC, before further incubation for 6 hrs when cells were then collected. Media was removed and cells were washed in PBS, typsinised and counted. For each sample 2 × 106 cells were pelleted in an Eppendorf, the supernatant was discarded and samples were frozen at -80 °C.

For analysis, samples were thawed and concentrated nitric acid was added (100 µL per sample) and mixed. Samples were then vortexed and sonicated and left overnight at room temperature. Samples were made up to 1 mL using Di water and then further diluted fivefold and analysed in the ICP Facility, School of Chemistry, University of Edinburgh.

### Nanostring Transcriptomic analysis

All NanoString nCounter analyses were carried out on the Human PanCancer Pathways (Catalogue number XT-CSO-PATH1-1) and Metabolic Pathways (Catalogue number XT-CSO-HMP-12) panels, covering 1449 genes.

Cells were seeded at 8 × 10^4^ cells in 6-well plates and incubated under standard tissue culture conditions for 24 hrs. For basal analysis media was then removed and plates were washed twice with ice cold PBS before being snap frozen at -80. For treatment induced analysis, media was replaced with fresh media containing compound treatments (DMSO (0.1%) or elesclomol (200 nM)) before further incubation for 6 hrs, and then samples were processed the same as for basal samples.

For RNA extraction cells were scraped and lysed using QIAshredders (#79654, QIAGEN), RNA was extracted by means of the Qiagen RNeasy Mini kit (#74104, QIAGEN) (with β-mercaptoethanol) according to manufacturer instructions, and included a DNase digestion step (#79254, QIAGEN).

## Results

We have explored NACs ability to alleviate the toxicity of the copper ionophores elesclomol and disulfiram, in two oesophageal cancer lines (OAC-P4C and SK-GT-4). We found, in line with previous research [8,12], that pretreatment with NAC alleviated elesclomol and disulfiram induced cell death (Fig 1).

**Fig 1.**
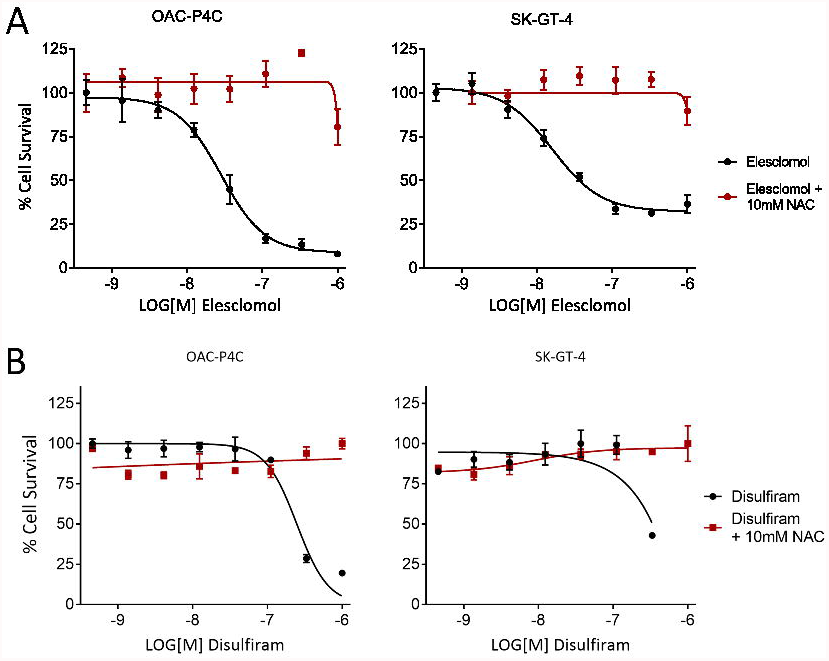
N-acetylcysteine prevents copper ionophore induced cell death at high concentrations. A) Elesclomol and B) Disulfiram dose response with and without 10mM N-acetylcysteine in OAC-P4C and SK-GT-4 cells. NAC= N-acetylcysteine

However, we found that NAC relieves elesclomol induced toxicity only at high concentrations (10mM). It has no effect at 1mM (Fig 2) which has been observed in other cell lines previously [12]. This finding provides evidence that NAC is not working via its antioxidant capacity, which is active at much lower concentrations [13], but rather is via alternative mechanisms requiring excess levels of NAC.

**Fig 2.**
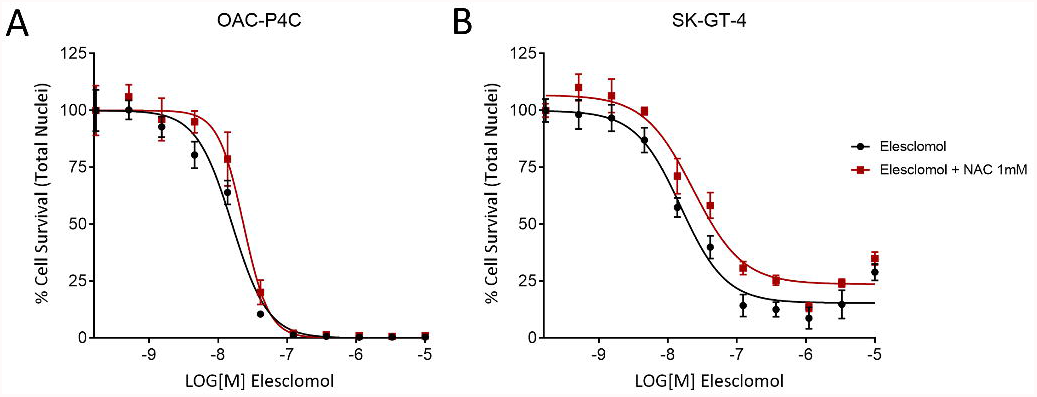
N-acetylcysteine does not abrogate elesclomol induced toxicity at 1mM. Elesclomol dose response in A) OAC-P4C and B) SK-GT-4 cells with and without 15 minute preincubation with 1mM N-acetylcysteine.

This observation is further supported by results demonstrating other antioxidants, for example trolox (data not shown) [14], and Coenzyme Q10 [12] do not alleviate the activity of elesclomol or disulfiram. Together, these results suggest NAC should not be used to demonstrate a role of ROS in copper ionophore mediated cell death.

Given these findings and the dual nature of NAC as an antioxidant and the less well known action as a copper interactor, we felt it was important to assess NACs effect on the intracellular copper levels since it could be acting via either mechanism. To our knowledge this has not been assessed previously. To quantify this, we measured the intracellular copper levels using inductively coupled plasma mass spectrometry (ICP-MS) in the OAC-P4C cell line. Importantly we demonstrate here for the first time that pre-incubation of cancer cells with NAC leads to complete loss of copper accumulation post elesclomol treatment, and results in copper levels comparable with those of the untreated cells (Fig 3). These results indicate that under these conditions NAC pretreatment dramatically blocks cellular uptake of copper via the copper ionophore elesclomol. Therefore, NAC pre-incubation appears to prevent compound activity by preventing the increase in intracellular copper levels and not via its antioxidant properties.

**Fig 3.**
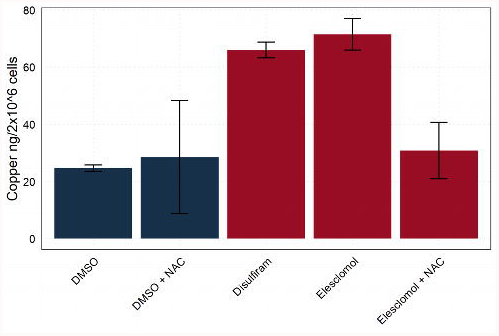
NAC prevents the intracellular increase in copper levels caused by copper ionophores. ICP-MS intracellular copper levels in the cell line OAC-P4C with and without 30 min, 10 mM NAC pre-treatment. NAC= N-acetylcysteine, ICP-MS=inductively coupled mass spectrometry.

To further investigate the mechanism of copper mediated cell death and any role of ROS, we explored elesclomol induced transcriptomic changes across a panel of oesophageal cancer cell lines using the NanoString nCounter platform. Firstly, assessment of the enzymes involved in maintenance of redox homeostasis did not reveal any major induction of antioxidant enzymes, including catalase (CAT) and superoxide dismutase (SOD) (Fig 4), suggesting the cells do not mount an antioxidant response. However, elesclomol treatment does lead to an increase in two genes involved in thiol homeostasis, TXNRD1 and GCLC, though this may be due to direct interaction of copper with thiol groups [15,16] rather than a result of ROS production.

**Fig 4.**
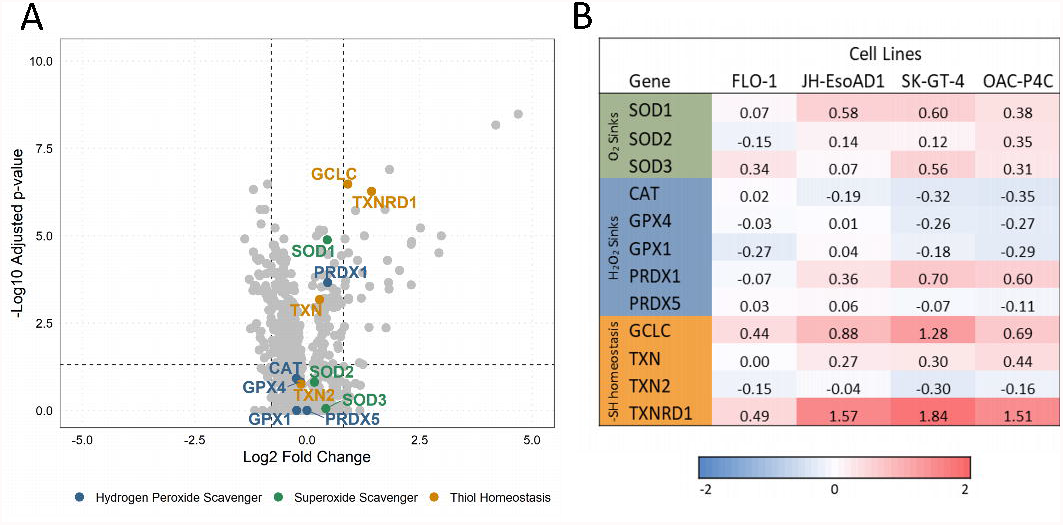
Elesclomol induced transcriptomic changes in redox homeostasis genes. A) Volcano plot for elesclomol induced differential expression analysis across FLO-1, JH-EsoAD1, SK-GT-4 and OAC-P4C cells. Redox homeostasis genes hydrogen peroxide scavengers (Blue), Superoxide scavengers (Green), Thiol homeostasis genes (Mustard) are labelled. B) Elesclomol induced transcriptomic log2 fold changes for highlighted redox homeostasis enzymes across four separate cell lines; FLO-1, JH-EsoAD1, SK-GT-4 and OAC-P4C cells.

Furthermore, we compared cell sensitivity to hydrogen peroxide (H_2_O_2_). If ROS levels are responsible for the cell death associated with the copper ionophores then we would expect the sensitivity to disulfiram and elesclomol to correlate with H_2_O_2_ sensitivity. We have previously profiled the sensitivity of 10 oesophageal cancer cell lines and two tissue matched controls to elesclomol and disulfiram and shown that the correlation in sensitivity to the two drugs is very high (0.94). Here we have taken the most sensitive cell line OAC-P4C with an IC50 of 1nM, and the most resistant cell line FLO-1, which is completely resistant even at concentrations of 10µM, and compared their sensitivity to H_2_O_2_. Results show that the copper ionophore resistant cells, FLO-1, are significantly more sensitive to oxidative stress via H_2_O_2_ than the copper ionophore sensitive cells, OAC-P4C, or tissue matched control cells (EPC2-hTERT) (Fig 5). This is consistent with previous findings that FLO-1 have a lower antioxidant capacity and increased sensitivity to ROS inducing compounds [17]. Overall, these findings do not fit with the hypothesis that ROS levels are responsible for copper mediated cell death, particularly in relation to the copper ionophores elesclomol and disulfiram.

**Fig 5.**
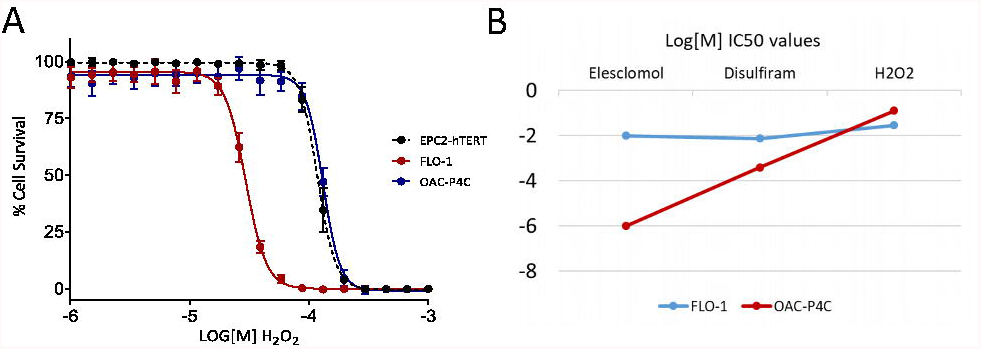
Sensitivity to copper ionophores and H_2_O_2_. A) H_2_O_2_ dose responses for OAC cell lines OAC-P4C and FLO-1, and tissue matched control EPC2-hTERT. B) Log molar IC50 values for Elesclomol, Disulfiram and H_2_O_2_, across the FLO-1 and OAC-P4C cell lines.

## Discussion

A new form of cell death termed cuproptosis has been discovered and reviewed extensively by others [18–20]. This new form of cell death has been widely studied in relation to a novel class of copper ionophores, including elesclomol and disulfiram. While there is a wealth of evidence that these compounds lead to the intracellular accumulation of copper, the exact mechanism leading to cell death remains contentious.

Since copper is a redox active metal, it has been proposed that the intracellular copper accumulation leads to the production of ROS and that this is the cause of cell death. Much of the evidence for this is based on the ability of the small molecule NAC to alleviate the toxicity of the drugs. Here we have demonstrated across multiple cell lines and drugs that NAC does indeed alleviate the cytotoxic effects of these copper ionophores when used at high millimolar concentrations. However, we have further shown that this is not through NAC’s canonical activity as an antioxidant but rather it significantly reduces copper cellular uptake, suggesting that reduced uptake of copper, rather than the antioxidant activity of NAC, is responsible for the diminished cell death. Given that NAC has and is still widely used in cell culture-based studies investigating the role of ROS generation in copper toxicity, we believe it is important to disseminate the finding that NAC interferes with copper ionophore mediated cellular uptake of copper, confounding conclusions drawn as to the effects of ROS.

We do not know the exact mechanism through which NAC significantly reduces copper cellular uptake caused by the ionophores but it may be either via direct copper sequestration/chelation or reduction of copper since NAC is a potent reducing agent and has been shown to potently reduce Cu(II) to Cu(I) [21] and this may prevent the elesclomol-copper chelate formation. Further disulfiram requires Cu(II) for its conversion to an active metabolite diethylthiocarbamate (DETC)[5] so in the presence of Cu(I) it will not undergo conversion to form copper chelates, potentially explaining its lack of activity.

Our results also demonstrate a lack of evidence that ROS is the major contributor to cuproptosis and copper ionophore induced cell death. Our transcriptomic results suggest that cells do not mount a strong antioxidant reaction in response to the copper ionophores. In fact, H_2_O_2_ sink enzymes glutathione peroxidase 1 and 4 (GPX1, GPX4), and catalase (CAT) are marginally downregulated. These copper ionophore mediated transcriptional changes mirror almost perfectly the results of a similar study on cuproptosis using excess concentrations of Cu (II) in the culture media [21] where they also demonstrate that there is no upregulation of antioxidant enzymes in response to excess copper. Collectively our data supports the more recent studies on cuproptosis that suggest while an accumulation of intracellular copper may well induce some levels of ROS, this is unlikely the mechanism leading to cuproptosis and cell death. Recent papers instead point towards protein aggregation and loss of proteostasis [1– 3,21].

Overall the results of this study indicate that NAC should be used with caution in experiments designed to ascertain the role of ROS in copper ionophore mediated cell death and cuproptosis in general and that this novel form of cell death requires further study to fully elucidate its mechanism.

## Acknowledgements

We would like to thank A. Rustig, University of Pennsylvania for provision of the EPC2-hTERT cells and a Cancer Research UK award (C42454/A24892) and the Anne Forrest Fund for Oesophageal Cancer Research for supporting our research.

## References

1. Hughes RE, Elliott RJR, Li X, Munro AF, Makda A, Carter RN, et al. Multiparametric High-Content Cell Painting Identifies Copper Ionophores as Selective Modulators of Esophageal Cancer Phenotypes. ACS Chem Biol. 2022;17: 1876–1889. doi:10.1021/acschembio.2c00301

2. Tsvetkov P, Detappe A, Cai K, Keys HR, Brune Z, Ying W, et al. Mitochondrial metabolism promotes adaptation to proteotoxic stress. Nat Chem Biol. 2019;15: 681–689. doi:10.1038/s41589-019-0291-9

3. Tsvetkov P, Coy S, Petrova B, Dreishpoon M, Verma A, Abdusamad M, et al. Copper induces cell death by targeting lipoylated TCA cycle proteins. Science. 2022;375: 1254–1261. doi:10.1126/science.abf0529

4. Nagai M, Vo NH, Shin Ogawa L, Chimmanamada D, Inoue T, Chu J, et al. The oncology drug elesclomol selectively transports copper to the mitochondria to induce oxidative stress in cancer cells. Free Radic Biol Med. 2012;52: 2142–2150. doi:10.1016/j.freeradbiomed.2012.03.017

5. Kragh HS. From Disulfiram to Antabuse: The Invention of a Drug. Bull Hist Chem. 2008;33: 82–88.

6. Dalecki AG, Haeili M, Shah S, Speer A, Niederweis M, Kutsch O, et al. Disulfiram and copper ions kill Mycobacterium tuberculosis in a synergistic manner. Antimicrob Agents Chemother. 2015;59: 4835–4844. doi:10.1128/aac.00692-15

7. Wu L, Zhou L, Liu DQ, Vogt FG, Kord AS. LC–MS/MS and density functional theory study of copper(II) and nickel(II) chelating complexes of elesclomol (a novel anticancer agent). J Pharm Biomed Anal. 2011;54: 331–336. doi:10.1016/j.jpba.2010.09.007

8. Kirshner JR, He S, Balasubramanyam V, Kepros J, Yang C-Y, Zhang M, et al. Elesclomol induces cancer cell apoptosis through oxidative stress. Mol Cancer Ther. 2008;7: 2319–2327. doi:10.1158/1535-7163.mct-08-0298

9. Barbi de Moura M, Vincent G, Fayewicz SL, Bateman NW, Hood BL, Sun M, et al. Mitochondrial Respiration - An Important Therapeutic Target in Melanoma. Santos J, editor. PLoS One. 2012;7: e40690. doi:10.1371/journal.pone.0040690

10. Zheng J, Lou JR, Zhang XX, Benbrook DM, Hanigan MH, Lind SE, et al. N-Acetylcysteine interacts with copper to generate hydrogen peroxide and selectively induce cancer cell death. Cancer Lett. 2010;298: 186–194. doi:10.1016/j.canlet.2010.07.003

11. Mi L, Sirajuddin P, Gan N, Wang X. A cautionary note on using N-acetylcysteine as an antagonist to assess isothiocyanate-induced reactive oxygen species-mediated apoptosis. Anal Biochem. 2010;405: 269–271. doi:10.1016/j.ab.2010.06.015

12. Buccarelli M, D’Alessandris QG, Matarrese P, Mollinari C, Signore M, Cappannini A, et al. Elesclomol-induced increase of mitochondrial reactive oxygen species impairs glioblastoma stem-like cell survival and tumor growth. J Exp Clin Cancer Res. 2021;40. doi:10.1186/S13046-021-02031-4

13. Zhang F, Lau SS, Monks TJ. The Cytoprotective Effect of N-acetyl-L-cysteine against ROS-Induced Cytotoxicity Is Independent of Its Ability to Enhance Glutathione Synthesis. Toxicological Sciences. 2011;120: 87–97. doi:10.1093/TOXSCI/KFQ364

14. Mittal M, Khan K, Pal S, Porwal K, China SP, Barbhuyan TK, et al. The Thiocarbamate Disulphide Drug, Disulfiram Induces Osteopenia in Rats by Inhibition of Osteoblast Function Due to Suppression of Acetaldehyde Dehydrogenase Activity. Toxicological Sciences. 2014;139: 257–270. doi:10.1093/TOXSCI/KFU020

15. Kreitman GY, Danilewicz JC, Jeffery DW, Elias RJ. Reaction Mechanisms of Metals with Hydrogen Sulfide and Thiols in Model Wine. Part 1: Copper-Catalyzed Oxidation. J Agric Food Chem. 2016;64: 4095–4104. doi:10.1021/ACS.JAFC.6B00641

16. Letelier ME, Lepe AM, Faúndez M, Salazar J, Marín R, Aracena P, et al. Possible mechanisms underlying copper-induced damage in biological membranes leading to cellular toxicity. Chem Biol Interact. 2005;151: 71–82. doi:10.1016/J.CBI.2004.12.004

17. Liu DS, Duong CP, Haupt S, Montgomery KG, House CM, Azar WJ, et al. Inhibiting the system xC- /glutathione axis selectively targets cancers with mutant-p53 accumulation. Nat Commun. 2017;8: 1–14. doi:10.1038/ncomms14844

18. Chen L, Min J, Wang F. Copper homeostasis and cuproptosis in health and disease. Signal Transduction and Targeted Therapy 2022 7:1. 2022;7: 1–16. doi:10.1038/s41392-022-01229-y

19. Tong X, Tang R, Xiao M, Xu J, Wang W, Zhang B, et al. Targeting cell death pathways for cancer therapy: recent developments in necroptosis, pyroptosis, ferroptosis, and cuproptosis research. Journal of Hematology & Oncology 2022 15:1. 2022;15: 1–32. doi:10.1186/S13045-022-01392-3

20. Xie J, Yang Y, Gao Y, He J. Cuproptosis: mechanisms and links with cancers. Molecular Cancer 2023 22:1. 2023;22: 1–30. doi:10.1186/S12943-023-01732-Y

21. Saporito-Magriñá CM, Musacco-Sebio RN, Andrieux G, Kook L, Orrego MT, Tuttolomondo MV, et al. Copper-induced cell death and the protective role of glutathione: The implication of impaired protein folding rather than oxidative stress. Metallomics. 2018;10: 1743–1754. doi:10.1039/c8mt00182k

